# Advancing Adoptive Cell Therapy: Optimized Expansion of Adaptive NK Cells for Solid Tumors

**DOI:** 10.1101/2024.10.02.616358

**Authors:** Nerea Martín Almazán, Sara Román, Yizhe Sun, Lars Bräutigam, Mattia Russel Pantalone, Giuseppe Stragliotto, Okan Gultekin, Sahar Saheli, Kaisa Lehti, Cecilia Söderberg Nauclér, Dhifaf Sarhan

**Affiliations:** Department of Laboratory Medicine, Division of Pathology, Karolinska Institute, Stockholm, Sweden; Department of Medicine, Solna, BioClinicum, Karolinska Institutet, 171 64, Stockholm, Sweden; Zebrafish Core Facility, Karolinska Institute, S-171 21, Stockholm, Sweden; Department of Neurology, Karolinska University Hospital, 171 76, Stockholm, Sweden; Department of Microbiology, Tumor and Cell Biology, Karolinska Institutet, Stockholm, Sweden; Department of Microbiology, Tumor and Cell Biology, Karolinska Institutet, Solnavägen 9, Stockholm 171 65, Sweden; Department of Women’s and Children’s Health, Division of Obstetrics and Gynecology, Karolinska Institutet, Stockholm, Sweden; Department of Pelvic Cancer, Theme Cancer, Karolinska University Hospital, Stockholm, Sweden; Department of Biomedical Laboratory Science, Norwegian University of Science and Technology, Trondheim, Norway; Department of Medicine Solna, Microbial Pathogenesis Unit, BioClinicum, Karolinska Institute, Stockholm, Sweden; Institute of BioMedicine, Unit of Infection and Immunology, MediCity Research Laboratory, Flagship InFLAMES, Turku University, Turku, Finland

**Author notes:** These authors contributed equally (Nerea Martín Almazán, Sara Román). Electronic address.

## Abstract

**Background:** Immune therapies are emerging as a critical component of cancer treatment, capable of delivering durable and potentially curative responses. While CAR-T cell therapy has proven effective for hematological malignancies, it faces challenges in treating solid tumors due to tumor antigen heterogeneity, an immunosuppressive tumor microenvironment, and physical barriers hindering CAR-T cell infiltration. NK cells, particularly adaptive NK (aNK) cells, offer a promising alternative due to their ability to recognize and kill tumor cells without prior sensitization and their resistance to immunosuppressive environments.

**Purpose:** The study investigates the role of cytokines, specifically IL-21 and IL-15, in enhancing aNK cell expansion and activation using peripheral blood mononuclear cells (PBMCs) from healthy donors and tumor-infiltrating lymphocytes (TILs) from glioblastoma (GBM) patients.

**Methods:** Buffy coats and GBM TILs were collected from Karolinska Hospital. NK cells were isolated and expanded in vitro with IL-15 and IL-21 cytokines and feeder cells (K562 and K562E). Furthermore, tumor lysate was added in the cultures to boost memory responses in aNK cells. NK cell functionality, cytotoxicity, and phenotyping was assessed using flow cytometry and statistical analysis (t-test and two-way ANOVA) used to validate the results. Further animal model was used to validate the cytotoxicity capacities of these cells against GBM tumors using a zebrafish model.

**Results:** IL-21 drives the expansion of aNK better than IL-15 controls, data shown in PBMCs and TILs derived from GBM patients and IPLA OVCA patients. Additionally, the use of tumor lysate as a booster for restimulation further amplifies the cytotoxic capacity of aNK cells against autologous tumors. The zebrafish model validates this method, by decreasing the tumor size in zebrafish animals after 3 days of injection.

**Conclusion:** The results demonstrate that IL-21 is essential for the specific expansion of aNK cells, enhancing their aggressiveness towards tumor cells. Additionally, tumor lysate significantly increases the cytotoxic efficacy of aNK cells upon restimulation with the same tumor cells. These findings suggest that IL-21 plays a crucial role in the specific expansion and activation of aNK cells, enhancing their aggressiveness towards tumor cells.

By optimizing the expansion protocol, this method aims to advance the clinical application of aNK cells in immunotherapies for solid tumors, offering a potential solution to the limitations faced by current CAR-T therapies.

## INTRODUCTION

Adoptive cell therapy (ACT), such as autologous chimeric antigen receptor (CAR) T cell therapy, has shown impressive results in the treatment of hematological malignancies, particularly in B-cell leukemia [1-4]. Unfortunately, this success cannot be applied to solid tumors, where CAR-T cells still have some challenges to overcome. These include the high tumor antigen heterogeneity, which makes it difficult for the CAR-T cells to detect cancer cells, the immunosuppressive tumor microenvironment (TME), where regulatory T cells (Tregs) and myeloid-derived suppressor cells (MDSCs) hamper the functionality and efficient killing of the effector cells, and physical and chemical barriers resulting in poor CAR-T infiltration into the tumor nest and consequently in tumor escape and the failure of tumor eradication [5, 6]. Thus, adoptive cell therapies require novel strategies for the treatment of solid malignancies.

Emerging evidence shows that Natural Killer (NK) cells with optimal anti-tumor properties may be a better option [7]. NK cells, as an important part of the innate immunity, are characterized by the expression of a complex array of cell-surface inhibitory and activating receptors and intracellular adaptor molecules that allow them not only to behave as significant regulators of immune responses but also, and more importantly, to develop strong cytotoxicity against virally infected and tumor cells without the need of prior sensitization [8-12]. According to the “missing self” hypothesis [13], NK cells’ inhibitory receptors, such as some of the Killer-cell Immunoglobulin-like Receptors (KIRs) and CD94-NKG2A heterodimer, recognize self-HLA class I molecules, dampening in this way the NK cell-mediated killing. However, the MHC class I downregulation in viral infections and cellular malignant transformation leads to imbalanced NK-target interactions that activate the NK cytotoxic pathways [11, 12]. In addition to this, despite their initial association with innate immunity, several studies have revealed the NK cell adaptive capacities, especially when they are exposed to cytomegalovirus (CMV) infections or cytokine treatment [14-19]. In fact, it is in this setting where a newly described subset of NK cells emerged in the field of immunology, the so-called “adaptive” NK (aNK) cells, due to their recently discovered properties: memory-like function and resistance to the immunosuppressor tumor microenvironment (TME) [20-22]. Different aNK cells derived from different contexts vary in terms of molecular signature [23, 24]. However, the most studied group are the aNK cells derived from CMV-infected individuals, whose memory capacities rely on the CMV-peptide presentation by the HLA-E on the stressed cell and its binding to the CD94-NKG2C heterodimer on the aNK cell [15, 25].

Considering the aforementioned properties of NK cells and aNK cells, it is apparent that NK cells attractively ascend to be an alternative in the immune cell-based immunotherapies in addition to T cells and therefore, *in vitro* or *ex vivo* methods to expand NK cells from multiple sources, including peripheral blood, are necessary to highly increase the NK cell number as well as reprogram the NK cell activity. In general, exposure to multiple cytokines potentiates NK cells to react towards the multiple ligands via binding to the assorted receptors [26]. IL-2 and IL-15 are considered primarily indispensable for NK cell development and proliferation [27, 28]. However, the outcomes from the clinical trials using IL-2 appeared unfavorable, owed to the side effects and insufficient specificity to NK cells, expanding Tregs instead [29, 30], and its combination with IL-15 failed to elicit the expected NK cell expansion [31]. Besides IL-2 and IL-15, the use of IL-21 as soluble or membrane-bound on feeder cell lines such as K562, K562-OX40L, or Epstein-Barr virus transformed lymphoblastoid cell line (EBV-LCL) has also been investigated [32-34], and recent results show that IL-21 is effective in cell therapy towards chronic lymphocytic leukemia (CLL), potentiating the NK cell expansion and elevating the tumor cytotoxicity [35].

As mentioned before, cytokine treatment has also been used to generate NK cell immune memory, and IL-12, IL-15, and IL-18 have demonstrated to be an efficient cytokine cocktail to induce sustained NK cell long-term responses in CMV-infected mice [18, 36]. In this context, the aNK cell subpopulation with adaptive features is arising in oncology clinical practices in relation to conventional NK cells (cNK) mainly due to its long-lasting specific responses and outperformed resistance to TME [20,21,24]. Consequently, and as well as for NK cells, the optimization of an aNK cell expansion protocol is required. In fact, feeder cells transgenetically modified to express HLA-E and mismatched HLA types seem to evolve as a key point for the selective expansion of human aNK cells [37-39]. Nevertheless, limited studies have so far been conducted to optimize the aNK cell expansion and further investigation for higher yields and enhanced functionality are still needed.

Alongside advancements in adaptive NK cell research, other therapeutic NK cell strategies— including cord-blood NK cells, stem cell-derived NK cells, cytokine-induced memory NK cells, and CAR-NK cells—are being investigated to improve NK cell-mediated therapies. The success of adaptive NK cell therapies, as well as these alternative approaches, depends on comprehensive evaluation in appropriate animal models to ensure both efficacy and safety [40].

Zebrafish (*Danio rerio*) embryos have emerged as a promising platform for high-throughput screening due to their small size, transparency, and cost-effective nature, which allow for high-throughput imaging of both cancer and immune cells. Notably, the lack of a fully functional adaptive immune system during early larval stages reduces the risk of cell rejection in xenograft studies [41]. Zebrafish xenografts have been utilized to investigate human tumor cell behaviors such as proliferation, apoptosis, invasion, and extravasation, and have been applied in anti-cancer drug screening. Recent studies have further demonstrated the zebrafish model’s ability to assess the in vivo efficacy of CAR-T cells [9, 42, 43]. These findings provide proof of concept that zebrafish xenografts can serve as a preclinical platform for testing CAR-T cell efficacy. However, NK cells differ from T cells in terms of proliferation, migration, activation, metabolic properties, and cytokine requirements. In a recent study by Murali Shankar et al. [41], the efficacy of CAR-NK cells against metastatic solid tumor cells was evaluated in a zebrafish model, successfully validating the CAR-NK system. They further examined the interactions between CAR-NK and cancer cells in real time, as well as the migration of CAR-NK cells through the microvasculature to metastatic breast tumors, demonstrating the zebrafish model’s value in preclinical cancer research.

In this study, we aim to investigate the combined role of IL-21 and IL-15 in promoting the expansion of adaptive NK (aNK) cells and to examine aNK cell activation in response to specific tumor antigens presented on HLA-E. Using peripheral blood mononuclear cells (PBMCs) from healthy donors as a model, with parallel validation in tumor-infiltrating lymphocytes (TILs), our goal is to optimize an *ex vivo* expansion protocol for aNK cells to facilitate their clinical application in adoptive NK cell-based immunotherapies. Additionally, we seek to assess the therapeutic potential of NK cell-mediated therapies by evaluating their efficacy using an appropriate animal model. To this end, we have developed an *in vivo* zebrafish model to study the tumor-killing activity of NK cells.

## MATERIALS AND METHODS

### Human participants

Blood from healthy donors (HD) was obtained from Karolinska Hospital (n=16). Two different tumor types were assessed: glioblastoma tumors collected from primary glioblastoma patients (n=6); and ovarian tumor samples collected from high-grade serous ovarian patients (n=8) participating in a Phase III clinical trial (Intra-Peritoneal Local Anesthetics in Ovarian Cancer trial (IPLA-OVCA), https://clinicaltrials.gov/ct2/show/NCT04065009). Written informed consents were obtained from all individuals before inclusion in the trial in accordance with the Declaration of Helsinki. The protocol for patient participation was approved by the local Ethics Committee and the Institutional Review Board (Dnº. 2019-05149 and 2015/1862-32).

### Sample processing

Peripheral blood mononuclear cells (PBMCs) were isolated from blood of HD by density gradient centrifugation using Ficoll-Paque Premium (GE Healthcare).

Tumor-infiltrating lymphocytes (TILs) were obtained from primary glioblastoma tumors or primary ovarian tumor biopsies, which were processed following a digestion method. The tumor specimen was cut in small pieces and incubated at 37ºC for 30 minutes in 5 mL of harvest media, consisting of RPMI1640 (Thermo Fisher Scientific) supplemented with 100 µg/mL RNase I (Sigma-Aldrich) and 150 µg/mL Liberase (Sigma-Aldrich). Following digestion, the solution was filtered through a strainer (Corning® cell strainer 70μm, Sigma-Aldrich) and washed in phosphate-buffered saline (PBS). Finally, the pellet was resuspended in 5 mL Red Blood Cell (RBC) lysis buffer, incubated at room temperature (RT) for 5 minutes, and washed in PBS. The final cell suspension was counted and divided into two fragments, one part for the TIL expansion and the other one for the expansion of primary glioblastoma tumor cells and primary ovarian tumor cells.

### Cell lines and primary tumor cells

Feeder cells K562 (A gift from Kärres group, original source) and K562E transfected with HLA-E*0101 (obtained from Dr. Michaëlesson’s laboratory, Karolinska Institutet) were grown in complete medium (CM), consisting of RPMI1640 supplemented with 10% heat-inactivated fetal bovine serum (FBS) (Thermo Fisher), 1% penicillin/streptomycin (100U/mL penicillin and 100 µg/mL streptomycin, Nordic Biolabs), and supplemented with 1mg/mL geneticin (Thermo Fisher). TYK-nu cells (Japanese Collection of Research Bioresources Cell Bank) were also cultured in CM. All cell lines were tested negative for mycoplasma contamination using the Mycoalert™ Mycoplasma Detection Kit (Lonza Bioscience Solutions) prior to use.

Human glioblastoma and ovarian tumor cells were expanded from the patient tumors and IPLA tumor biopsies, and the tumor cell suspension was cultured in a T-75 flask with CM supplemented for the glioblatoma tumor cells. In addition to CM, ovarian tumor cells with 10% autologous patient-derived ascites and 1 µg/mL ciprofloxacin hydrochloride (GenHunter). The outgrowth of cells was monitored on a weekly basis. In order to obtain a primary tumor cell culture free of tumor-associated cells, such as lymphocyte subpopulations, adipocytes, fibroblasts or endothelial cells, once 90% confluency was reached, the tumor cells were harvested and isolated using the Tumor Cell Isolation Kit (Miltenyi Biotec) following the manufacturer’s instructions. Tumor cell purity was assessed though the expression of the tumor/epithelial marker pan Cytokeratin (PanCK) (Abcam) and the lack of the fibroblast marker S100A4 expression (Lifespan Biosciences).

### Tumor cell lysate preparation

The UL18 glioblastoma cell line and TYK-nu ovarian cancer cell line was harvested, washed with PBS, and resuspended at 10^7^ cells/mL in PBS. Cells were lysed after six successive freeze-thawing cycles in liquid nitrogen and a 56ºC water bath. The resulting lysate, UL18 lysate or TYK-nu lysate, was filtered using a 70 µM cell strainer. Lastly, the protein concentration was measured using a NanoDropTM Protein A280 (Thermo Fischer Scientific), and the lysates were stored at −20ºC until use.

### *Ex vivo* expansion of aNK cells from PBMCs or TILs

PBMCs/TILs were co-cultured with 100 Gy irradiated K562 or K562E at 5:1 ratio (feeder cells : PBMCs/TILs) in 24-well plates at 0,5M PBMCs/TILs / well. Cells were cultured in RPMI1640 supplemented with 10% human AB-serum (Stockholm Blood Center) and 1% penicillin/streptomycin for 21 days, and 100 Gy irradiated K562E feeder cells were readded at 1:1 – 5:1 ratio on days 7 and 14. Some of the cultures were also supplemented with 5 ng/mL of exogenous cytokines (Miltenyi Biotec) recombinant human rh-IL15 or rh-IL15+rh-IL21, on days 0, 3, 7, 10, 14, and 17, and the medium was exchanged every 3-4 days. When necessary, the cell suspensions were transferred to 12-well plates, 6-well plates, T-25 flasks, T-75 flasks or T-175 flasks to always maintain the cell density at 0,5-1M total cells/mL. Alternatively, prior to irradiation, feeder cells were incubated O.N. with 3 µg/M cells of UL18 or TYK-nu lysate or 10 µg/mL of HLA-G-leader-derived-peptide (VMAPRTLFL, Nordicbiosite) according to previously published aNK cell expansion protocol (39) as a control.

### Expansion monitoring of the immune cells by flow cytometry

To monitor the expansion of the different subsets of immune cells, cells were weekly stained with anti-human CD3 (BV510, BD Biosciences), CD56 (PE-Cy7, BioLegend), CD57 (APC, BioLegend), NKG2C (PE, BioLegend) antibodies, then fixed and permeabilized (eBioscienceTM FOXP3/Transcription Factor Fixation/Permeabilization buffer, Thermo Fisher Scientific) for 20 min at RT in the dark to be intracellularly stained with anti-human FCεRIγ (FITC, Milli-Mark) and SYK (FITC, BioSciences). Here, we also used LIVE/DEAD Fixable Near-IR Dead Cell Stain Kit (Thermo Fisher Scientific) as live/dead cell fixable dye (APC-Cy7), as well as anti-human CD45 (BV421, BD Biosciences) for the TILs expansion. All samples were acquired by CytoFLEX cytometer (Beckman Coulter Life Sciences) and FlowJo v.10.8.1 software (BD Biosciences) was used to quantify the frequency of NK cells (CD3-CD56+), T cells (CD3+ CD56-), NKT cells (CD3+ CD56+), aNK cells (CD3-CD56+CD57+ Fc□RI□-, NKG2C+) and cNK cells (CD3-CD56+CD57+ FcεRIγ+, NKG2C-) in each culture.

### Functional assay by flow cytometry

The functionality of the aNK and cNK cells was assessed for degranulation (CD107a) and pro-inflammatory cytokine production IFNγ and TNFα, by general restimulation (days 7, 14, and 21) and specific restimulation (days 14, and 21) of the PBMCs/TILs. The general restimulation was performed with 50 ng/mL phorbol myristate acetate (PMA) (Sigma-Aldrich) and 1µg/mL ionomycin (Sigma-Aldrich), and the specific restimulation with viable tumor cells (UL18 and/or GBM autologous primary tumor cells or TYK-nu or/and autologous primary ovarian tumor cells) at 5:1 E:T ratio. In both cases, restimulation was undertaken in RPMI1640 medium together with anti-human CD107a antibody (PerCP/Cy5.5, BioLegend), protein transport inhibitors Golgiplug (BD Biosciences) (1:1000) and Golgistop (BD Biosciences) (1:1000) for 6h incubation prior to staining. Following surface staining, cells were fixed and permeabilized for 20 min at RT in the dark, and later intracellularly stained with anti-human intracellular FCεRγI, SYK, IFNγ (BV650, BD Biosciences) and TNFα (BV785, BioLegend). All samples were acquired by CytoFLEX cytometer and FlowJo v.10.8.1 software was used for the analysis.

### Phenotyping of the aNK and cNK cells from PBMCs and TILs by flow cytometry

To follow-up the aNK and cNK cells during the three weeks of expansion, their phenotype was assessed on days −1 and 21. Cells were stained with live/dead cell fixable dye and for different NK cell receptors – CD45 (PE-Cy7, BD BioSciences) CD3 (APC-Cy7, BioLegend), CD56, CD57 (BV605, BioLegend), NKG2C, NKG2A (APC, R&D Systems), NKG2D (AF700, R&D Systems), NKp30 (PerCP/Cy5.5, BioLegend), NKp44 (BV510, BD BioSciences), DNAM1 (BV650, BD BioSciences), TIM3 (BV421, BioLegend), and TIGIT (BV785, BioLegend) – and then fixed and permeabilized for 20 min at RT in the dark to be intracellularly stained with anti-human FCεRIγ and SYK. All samples were acquired by CytoFLEX cytometer and FlowJo v.10.8.1 software was used for the analysis.

### Killing assay by flow cytometry

To examine the killing capacities of the expanded aNK cells against autologous and/or allogeneic tumor cells at the end of the expansion (day 21), PBMCs and TILs were co-cultured during 6h with viable tumor cells (UL18 cells or TYK-nu cells and/or autologous primary glioblastoma or ovarian tumor cells), marked with CellTracker Green CMFDA (FITC, Thermo Fisher Scientific), at 1:1 and 5:1 E:T ratio. After incubation, cells were stained with live/dead cell fixable dye and anti-human CD45 (BV421, BD Biosciences), CD3 (BV785, BD BioSciences), and CD56 (PE, BioLegend) antibodies, and then fixed with 4% paraformaldehyde. All samples were acquired by CytoFLEX cytometer and FlowJo v.10.8.1 software was used for the analysis.

### Zebrafish model

According to the EU directive 2010/63/EU and the Swedish legislation on animal experimentation (L150), zebrafish embryos younger than 5 days are not considered research animals. Breeding animals are kept under the ethical license #15591/2023. The facility registration number for animal experimentation is: #5.2.18-05581/2020 The facility registration number for keeping aquatic animals is: #1142.

Zebrafish were housed in self-cleaning 3.5 l tanks with a density of 5 fish per liter in a centralized recirculatory aquatic system (Tecniplast). Basic water parameters were continuously surveilled and automatically adjusted to a temperature of 28 °C; conductivity 1200 µS/cm, pH 7.5. Other chemical water parameters were checked minimum monthly. The lightning scheme was 14 hours light/10 hours dark with a 20 min dawn and dusk period.

Any animals are imported to a physically separate quarantine unit from which only surface-disinfected eggs are transferred to the breeding colony barrier. Health monitoring was done through Charles River according to the FELASA-AALAS guidelines (PMID: 35513000). Mycobacterium chelonae has been detected in randomly sampled fish and in sludge samples, Mycobacterium fortuitum has been detected in sludge samples. ZfPV-1 has been detected in sentinel fish, randomly sampled fish and sludge. Zebrafish embryos were staged according to Kimmel et al. (PMID: 8589427). All husbandry procedures are defined in SOPs which are available together with the latest health monitoring reports on request.

Embryos are incubated in E3 medium at 28.5 °C until they reach the 1k-cell stage (4 hours post fertilization). Directly before transplantation, UL18 cells are spun down (2000 rpm for 90 sec), almost all buffer is removed, and the pellet is resuspended in a minimal volume of remaining buffer by flipping the tube. A microcapillary (TW100-4, World Precision Instruments) is pulled using a Sutter P1000 needle puller. The concentrated cell suspension is loaded into the microcapillary which is then connected to a Femtojet 4x (Eppendorf). The tip of the needle is opened with a blade. Just before transplanting, the embryos are lined up on an injection plate (petri dish covered with 1 % agarose prepared in E3 medium on which a mold has been placed which forms grooves into which the embryos can be placed). About 100-300 UL18 expressing mCherry cells are injected into the center of the cell mass. After transplantation, the embryos are collected in E3 medium and raised at 33°C.

Zebrafish embryos are raised to 48 hours post fertilization (hpf) in E3 water containing 30 mg/l phenylthiourea (PTU) to block pigmentation. PTU is added to the embryos at 4 hours post fertilization (hpf). Directly before transplantation, cultures of long-term PBMCs cultured with different cytokine cocktails and K562 or K562-E cells are spun down (2000 rpm for 90 sec), almost all buffer is removed, and the cell pellet is resuspended in a minimal volume of remaining buffer by flipping the tube. A microcapillary (TW100-4, World Precision Instruments) is pulled using a Sutter P1000 needle puller. The concentrated cell suspension is loaded into the microcapillary which is then connected to a Femtojet 4x (Eppendorf). The tip of the needle is opened with a blade. Just before transplanting, the embryos are anesthetized using 160 µg/ml Tricaine (Pharmaq AS #434261). Embryos are transferred onto an agarose surface and placed on their ventral side. About 100-300 cells are injected into the common cardinal vein (Duct of Cuvier). After transplantation, embryos are screened for successful transplantation and incubated at 33°C.

The imaging 96-well plate (Ibidi, #89621) is prepared in the following way: 1% agarose is prepared in 1xE3 medium and 250 µl are distributed per well. A mold (prepared by the 3D-printing core facility at Uppsala University based on PMID:24886511) is placed on top of the plate until the agarose has solidified. Using a 1000 µl pipette, single embryos are distributed in 150 µl exposure medium (160 µg/ml tricaine, 30 mg/l phenylthiourea in E3 medium) distributed into the wells of the 96-well imaging plate. Embryos are manually oriented into position and imaged as described below. The embryos are kept at 33°C during the imaging and moved to a wet-chamber and 33°C incubator after imaging.

Confocal imaging is performed using brightfield (4x) and red channel to detect the signal from the UL18 mCherry cells.

### Confocal microscopy

Zebrafish animals were sectioned at 3μm in the histology lab at KI. These sections were rehydrated in increasing percentages of water (xylene, 100% ethanol, 95% ethanol, 70% ethanols and PBS) and a citrate-based solution (Vector Antigen Unmasking Solution Cat H3300) was done for antigen retrieval. Samples were permeabilized with 0,1% Triton X-100 and blocked with PBS containing 5% of goat serum. After blocking, samples were stained with CD45 (FITC, BD Biosciences) and Hoersch antibody (ThermoFisher Scientific). Samples were mounted and observed by confocal microscopy Zeiss.

### Bioinformatic analysis

The online tool GEPIA 2 (www.gepia2.cancer-pku.cn) was used to perform a correlation analysis between the aNK cell’ signature and the genes encoding for IL-15, IL-21, and HLA-E, using Pearson coefficient. The expression dataset (glioblastoma or ovarian cancer) was collected from the public database TCGA.

### Statistical analysis

All experiments were repeated independently at least three times. One representative and accumulative data are presented. Before further statistical analysis, all numeric data were subjected to normal distribution using the Shapiro-Wilk test and QQ plots tests. For the comparison within groups, either a multiple paired T test without considering the effects within groups or parametric two-way ANOVA we performed. Student T-test was used when comparing two groups. Correlation analyses (Pearson coefficient) were performed. All statistical tests were two-sided. All p-values from multiple comparisons were corrected by using the FDR method <0.05. The Prism v9.2 software (GraphPad) was used for statistical analyses.

## RESULTS

### Efficient expansion of adaptive NK cells (aNK) using IL-15+IL-21 cytokines in PBMCs

The selective expansion of aNK cells has already been optimized using IL-2 and IL-15 (37– 39); however, here we would like to describe a new expansion protocol (**Fig. 1a**) using a different cytokine cocktail, IL-15+IL-21, along with K562E feeder cells, since IL-21 has demonstrated to enhance both the expansion and tumor cytotoxicity of human NK cells (32– 34). Furthermore, the correlation analysis between the aNK cell gene signature and the genes encoding for IL-15, IL-21, and HLA-E has a lower p-value and a higher R-value than comparing the aNK cells’ signature to IL-15 (p-value=0,02; R=0,002) or IL-15 and IL-21 (p-value=1,7e-06; R=0,23) (**Fig. 1b**), which supports the feasibility of the expansion protocol. For its optimization, three different conditions were tested HD-derived PBMCs: K562E, K562E+IL-15, and K562E+IL-15+IL-21. We found that stimulation with exogenous cytokines potentiates the expansion of NK cells, and especially the cytokine cocktail of IL-15+IL-21, rather than IL-15 alone (**Fig. 1c, 1d**). At the end of the expansion, NK cells reach a mean cell frequency of 80% without the need for prior bead-isolation from bulk PBMCs, and no T cell contamination was found in the combination of IL-15+IL-21, however, residual monocytes may still be in culture (**Fig. 1c, 1d**). Within the NK cells, we also demonstrated that the cytokine cocktail of IL-15+IL-21 enhances the selective expansion of aNK cells over cNK cells without prior bead-isolation (**Fig. 1e, 1f**), allowing us to obtain a mean of 450-expansion fold of aNK cells (**Fig. 1g**) without prior bead-isolation. The fold change of aNK cell count was around 16 times at day 14 compared to day 7, and it decreased to 2 at day 21 compared to day 14. In cNK cells, we did not observe such an increase in any of the days (**Fig. 1g**). In addition, in IPLA TILs derived from ovarian cancer, we observed an NK cell median of 14000-expansion of of 14000-fold was reached (**Fig. 1h**). We also noticed that the proliferation rate of the aNK cells decreases along the expansion (**Fig.1h**), which might be explained by the lack of antigen stimulation.

**Figure 1.**
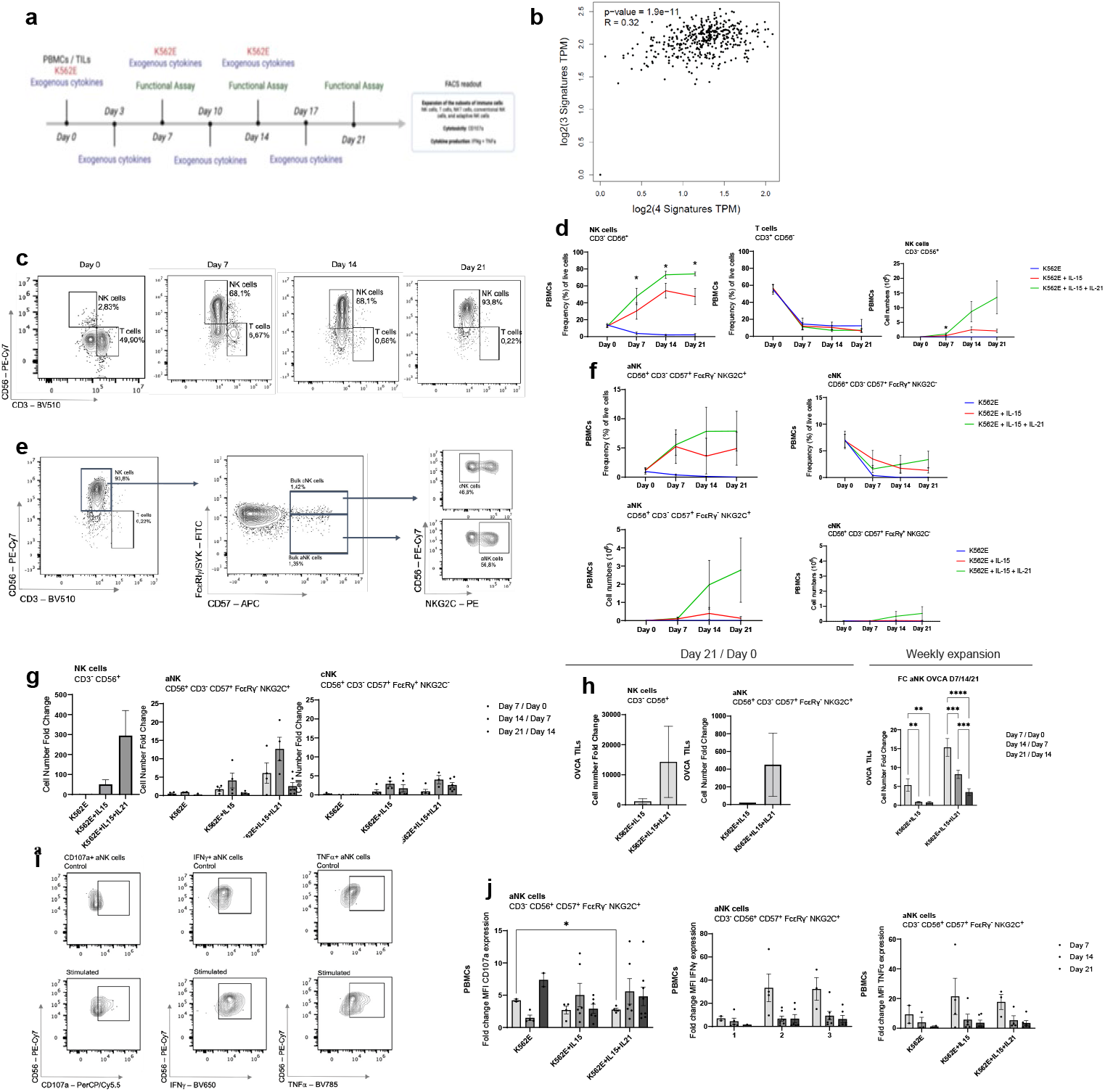
Expansion analysis of adaptive NK cells using K52E feeder cells with exogenous cytokines IL-15 or IL-15+IL-21. **a**. Protocol procedure for the selective expansion of aNK cells. Figure created with BioRender.com. **b**. Correlation analysis (Pearson) between *NCAM1, B3GAT1, KLRC2*, and *CD3E* (aNK cells’gene signature) and *IL15, IL21* and *HLA-E* performed on a TCGA expression dataset from ovarian cancer patients (left) and GBM patients (right) (www.gepia2.cancer-pku.cn) **c**. Expansion analysis by flow cytometry of NK cells and T cells on days 0-21. Gating was performed on viabled CD45 ^+^ cells and subsequently on CD3 ^-^/CD56^+^ for NK cells and CD3 ^+^/CD56^-^ for T cells. **d**. Frequency of NK and T cells of live cells as well as cell numbers of NK cells during expansion. **e**. Gating strategy on a representative HD for determining the expansion of aNK cells. **i**. Gating was performed on viable CD45^+^ cells and subsequently on CD3 ^-^, CD56 ^+^, CD57 ^+^, FcεRγ^-^, and NKG2C+ NK cells. **f**. Frequency of live cells and cell numbers of aNK and cNK cells during the expansion. **g**. Fold change of aNK cells and cNK cell numbers after expansion. **h**. Fold change of NK cells and aNK cells derived from ovarian cancer tI TILs **i**. Gatingting strategy on a representative PBMCs donor for analyzing the functionality of aNK cells in terms of degranulation (CD107a) and cytokine production (IFNγ and TNFα). Functional assay by flow cytometry performed on aNK cells on days 7, 14, and 21 of the expansion. The degranulation and cytokine production are studied with the MFI fold change, meaning MFI (stimulated sample) / MFI (unstimulated smaple). **j**. MFI Fold Change of aNK cells on PBMCs degranulation and cytokine production.

Upon general stimulation, aNK cells have augmented cytotoxicity and cytokine production capacities (**Fig. 1i**), especially when they are expanded under biweekly stimulation with exogenous cytokines; however, no significant differences were found in the functionality of aNK cells between their expansion with IL-15 alone or IL-15+IL-21, suggesting that IL-21 has higher impact on the expansion of aNK cells and maintains their functional capacity (**Fig. 1j**). aNK cells time-dependently increased in degranulation upon exogenous IL-15 cytokine stimulation (**Fig. 1j**). In contrast, the highest activity for IFNγ and TNFα production was reached during the first week of expansion and back to baseline following long-term stimulation (**Fig. 1j**). These results suggest that aNK cells may need antigen-specific stimulation that mainly drives their activation.

### Efficient expansion of adaptive NK cells (aNK) using IL-15+IL-21 cytokines in TILs

After observing these results on PBMCs, we validated the expansion of NK cells and aNK cells on TILs derived from GBM patients and ovarian cancer patients. We used the three different conditions of the cytokine cocktail and incubated them with K562E. Results with K562 can be found in supplementary figure 1. As observed in PBMCs, the conditions that contained IL-21 expanded better NK cells (**Fig. 2a**), in both cocktails T cells frequencies decreased. Regarding aNK cells, IL-21 controls expanded better aNK cells (**Fig. 2b**) and cNK cells decreased in both controls.

**Figure 2.**
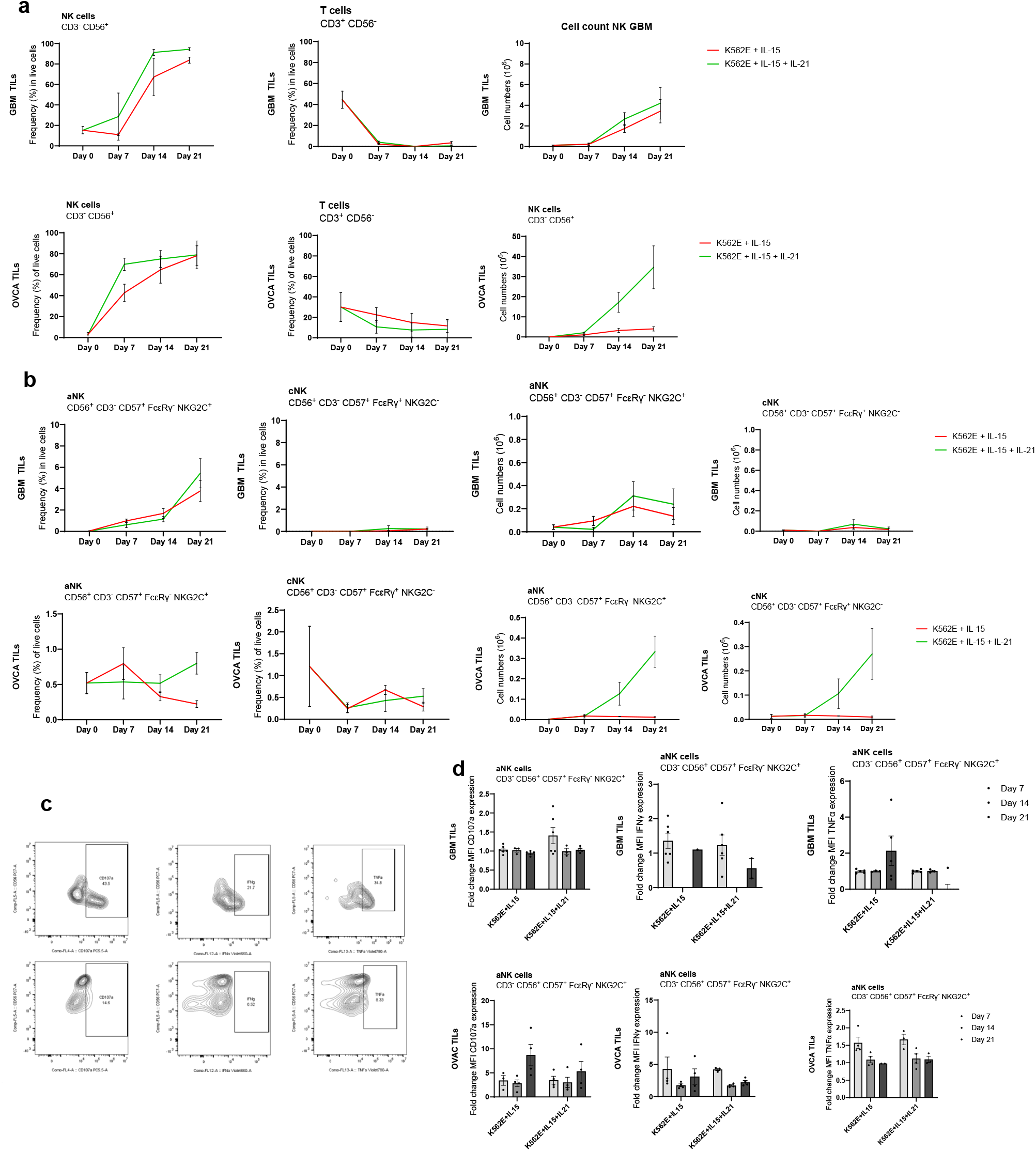
Expansion analysis of adaptive NK cells using K52E feeder cells with exogenous cytokines IL-15 or IL-15+IL-21. **a**. Frequency of NK and T cells of live cells as well as cell numbers of NK cells during expansion. **b**. Frequency of live cells and cell numbers of aNK and cNK cells during the expansion. **c**. Functionality of aNK cells during long-term expansion in order to characterize the role of aNK cells. Gating strategy of a representative GBM donor for analyzing the functionality of aNK cells in terms of degranulation (CD107a) and cytokine production (IFNγ and TNFα). Functional assay by flow cytometry performed on aNK cells on days 7, 14 and 21 of the expansion. The degranulation and cytokine production are studied with the MFI fold change, meaning MFI (stimulated sample) / MFI (unstimulated sample).

Upon general stimulation, aNK cells derived from GBM TILs or ovarian cancer TILs have augmented cytotoxicity and cytokine production capacities (**Fig. 2c, d**), especially when they are expanded under biweekly stimulation with exogenous cytokines; however, no significant differences were found in the functionality of aNK cells between their expansion with IL-15 alone or IL-15+IL-21, suggesting that IL-21 has higher impact on the expansion of aNK cells and maintains their functional capacity (**Fig. 2d**). aNK cells time-dependently increased in degranulation upon exogenous IL-15 cytokine stimulation (**Fig. 2d**). In contrast, the highest activity for IFN γ and TNFα production was reached during the first week of expansion and back to baseline following long-term stimulation (**Fig. 2d**). These results suggest that aNK cells may need antigen specific stimulation that mainly drives their activation.

### IL-21 maintains natural aNK cells features

Next, we showed that the expansion protocol with IL-21 does not negatively affect the expression of the NK cell activating receptors, but rather it successfully maintains the natural aNK cells’ features (**Fig. 3a, b**).

**Figure 3.**
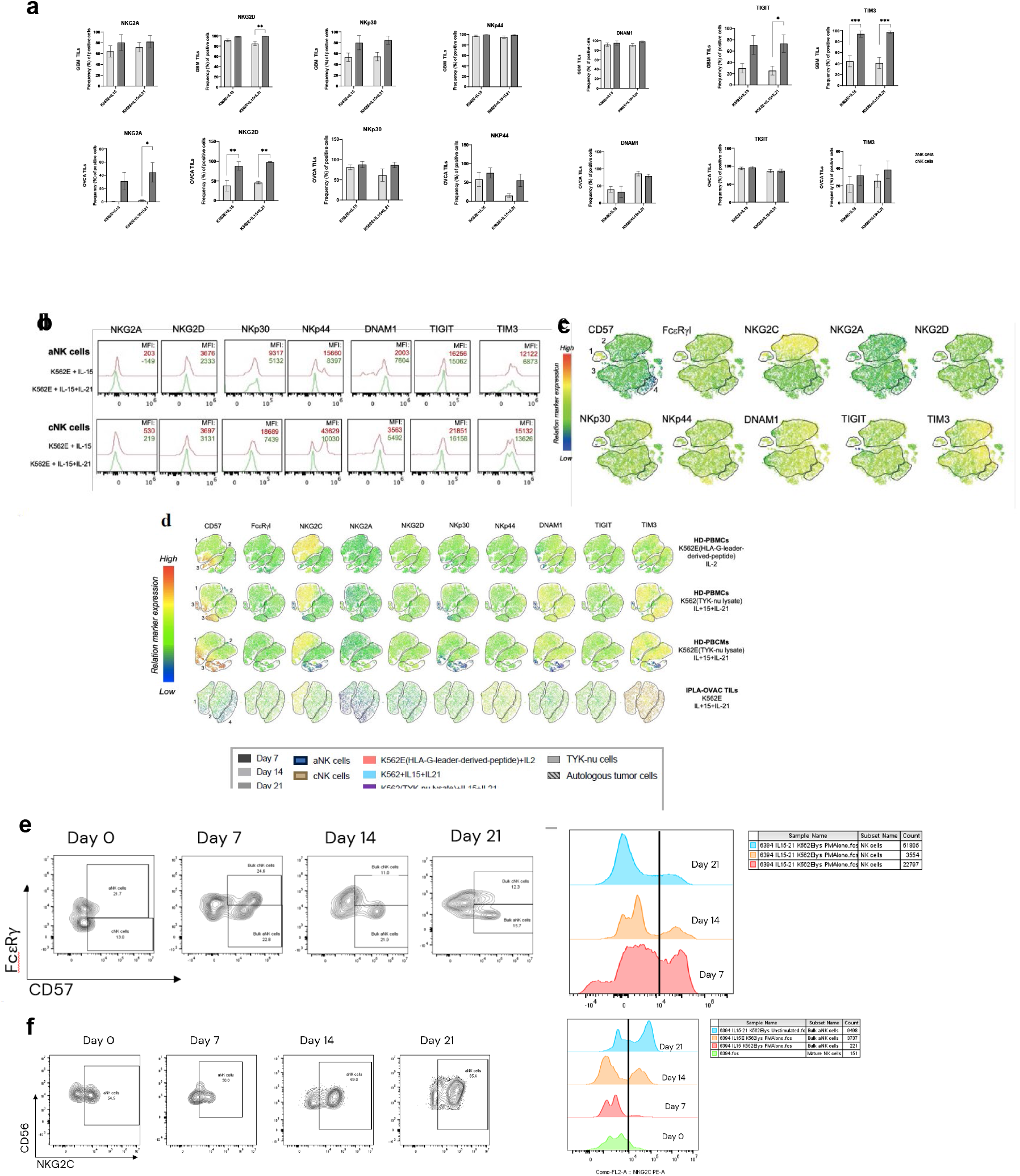
Phenotyping of aNK cells during long-term expansion in order to characterize the role of IL-21. **a**. Differential expression of NK receptors on aNK and cNK cells on day 21 in terms of cell frequency and MFI. The phenotyping was performed on TILs (*n=3*), and the histograms show the receptors’expression of NK receptors from a representative IPLA-OVCA donor. **b**. Relative expression of bulk NK cells for the assessed markers by flow cytometry shown on a t-SNE clustering plot. The presented data is from a representative IPLA-OVCA donor on day 21 under the condition of K562E + IL-15 + IL-21, and four different clusters are identified: (1,2) aNK cells, (3) intermediate aNK cells, and (4) immature NK cells. The positivity of the functionality and phenotyping markers was determined with Fluorescence Minus One (FMO) tests. Data is presented as mean ± standard error of the mean (SEM), and statistical differences were tested using and two-way ANOVA; *p<0,05, **p<0,01. **c**. t-SNE plots indicating the relative expression of the markers shown in b. **d**. t-SNE plots of the same markers from NKcells stimulated on different conditions between HD and TILs. **e**. FACS plots showing the maturation (CD57+ at days 0, 7, 14 and 21. **f**. FACS plots showing the NKG2C+ expression at days 0, 7, 14 and 21.

Within the family of CD94/NKG2 receptors, NKG2A is an inhibitory receptor that competes with NKG2C, an activating receptor, for the binding to HLA-E and, as it was previously demonstrated by Almazán et al. its downregulation on aNK cells, compared to its upregulation on cNK cells, is key for the aNK cell-activation against target cells (**Fig. 3a, b, c**). NKG2D, on the contrary, is an activating NK receptor that is expressed on both aNK and cNK cells, which is essential for the NK-killing (Billadeau DD et al. Nature Immunol 2003), however it is lower on aNK cells (**Fig. 3a, b, c**). Regarding the Natural Cytotoxicity Receptors (NCRs), here we show that both NKp30 and NKp44 are downregulated in aNK cells compared to cNK cells (**Fig. 3a, b**), especially when IL-21 is part of the cytokine cocktail, which can be explained by the epigenetic changes aNK cells go through to develop memory-like function. In regard of the expression of DNAM1, TIGIT, and TIM3, DNAM1 and TIM3 are activating receptors (Pazina et al. Cancers, 2021) that are highly expressed on both aNK and cNK cells, especially with the use of IL-21 (**Fig. 3a, b**), unlike TIGIT, which is an inhibitory receptor that, as has been previously demonstrated by Sarhan et al. (Sarhan et al. Cancer Res. 2016), is downregulated on aNK cells compared to cNK cells.

NKp44 has been reported as an activating NCR, especially under the expansion with IL-15, enhancing the cytotoxicity and IFNγ and TNFα production of NK cells (Sivori S. et al., Cell Mol Immunol 2019). However, here we show that the expansion with IL-15+IL-21 highly correlates the NKp44-expression on aNK cells with their killing and immunoregulating capacities, which is the opposite to the expansion with IL-15 alone (**Fig. 3d**), implying the IL-21 potentiates the activated phenotyping of the aNK cells.

During the expansion of aNK cells, we observed that NK cells became more mature expressing higher CD57 (**Fig. 3e**) and they expressed higher NKG2C at day 21 compared to day 0 (**Fig. 3f**).

### Efficient expansion of aNK cells in the presence of IL-21 and tumor lysate

The selective expansion of aNK cells using IL-15 and IL-21 has already been discussed in Figures 1, and 3; however, here we would like to describe a new expansion protocol (**Fig. 4a**) using the same cytokine cocktail, IL-15+IL-21, along with K562E feeder cells, and adding tumor lysate into the cultures. Lysates derived from UL18 GBM cell line and TYK-nu ovarian cancer cell line were used to expand aNK cells and study their role in memory responses against tumor cells. For its optimization, three different conditions were tested HD-derived PBMCs and TILs (GBM and ovarian cancer): K562E, K562E+IL-15, and K562E+IL-15+IL-21. We found that stimulation with exogenous cytokines and tumor lysate potentiates the expansion of NK cells, similar to cultures with no lysate. This expansion is particularly in the conditions with IL-15 + IL-21 (**Fig. 4b**). At the end of the expansion, NK cells reach a mean cell frequency of 80% without the need of prior bead-isolation from bulk PBMCs, and no T cell contamination was found in the combination of IL-15+IL-21, however, residual monocytes may still be in culture (**Fig. 4b, 4c**). Within the NK cells, we also demonstrated that the cytokine cocktail of IL-15+IL-21 enhances the selective expansion of aNK cells over cNK cells without prior bead-isolation (**4d**). In previous cultures with no antigen stimulation, the population of aNK cells decreased overtime. In these cultures with lysate, we observed that the aNK cell did not increase overtime but kept expanding (**Fig. 4d**). Mention results of HlA-G leader peptide.

**Figure 4.**
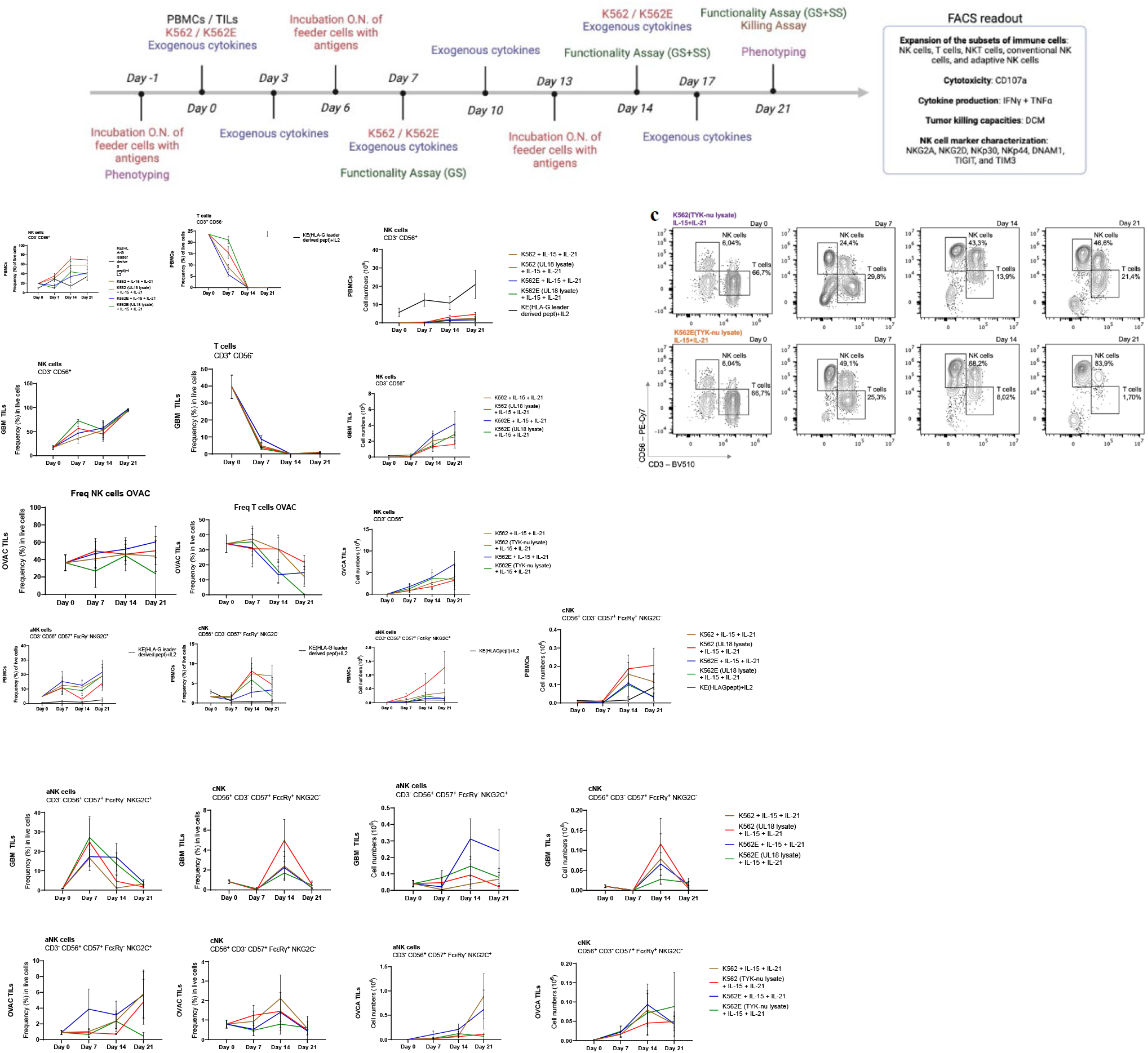
Expansion analysis of NK cells and adaptive NK cells upon tumor antigen stimulation. **a**. Protocol procedure for the selective expansion of aNK cells. Figure created with BioRender.com. **b**. Frequency of NK and T cells of live cells as well as cell numbers of NK cells during the expansion. **c**. Expansion analysis by flow cytometry of NK cells and T cells on days 0-21 from a representative HD. Gating was performed on viable CD45^+^ cells and subsequently on CD3^-^CD56^+^ for NK cells and CD3 ^+^/CD56^-^ for T cells. **d**. Frequency of live cells and cell numbers of aNK and cNK cells furing the expansion. In all cases, HD-derived PBMCs (*n=9*), GBM TILs (*n=6*) and IPLA-OVAC TILs (*n=4*) were used as samples, and cells were expanded under IL-15+IL-21 stimulation together with K562/K562E feeder cells with or without overnight exposure to UL18 lysate (for GBM TILs) or TYK-nu lysate (for OVAC TILs). Data is presented as mean ± standard error of the mean (SEM), and statistical differences were rested using multiple paired t-tests, paired t-tests when comparing two groups, and multiple Wilcoxon test when analyzing the frequency and cell number of aNK cells (not normally distributed); *p<0,05, **p<0,01.

### IL-21 and tumor lysate maintain natural aNK cells’ features with high functionality along long-term expansion

As suggested in Fig 1, aNK cells need specific stimulation to drive their activation and functionality. As hypothesized, the degranulation (measured by CD107a expression) and cytokine release (measured by IFNγ and TNFα expression) increased over time in HD cultures that were exposed to tumor lysate, compared to the cultures that were not (**Fig 5a, 5b**). This result is particularly opposite to the HLA-G leader peptide control, which decreases overtime in CD107a or is unchanged in cytokine release. In TILs cultures, we did not observed differences in CD107a or cytokine release with general stimulation, but when these TILs were cultured with tumor cells (autologous or UL18 cell line or TYK-nu cell line) the cytotoxicity of these aNK cells was higher when stimulated with autologous tumor cells compared to stimulation with a general cell line derived from GBM or ovarian cancer (**Fig. 5d**).

**Figure 5.**
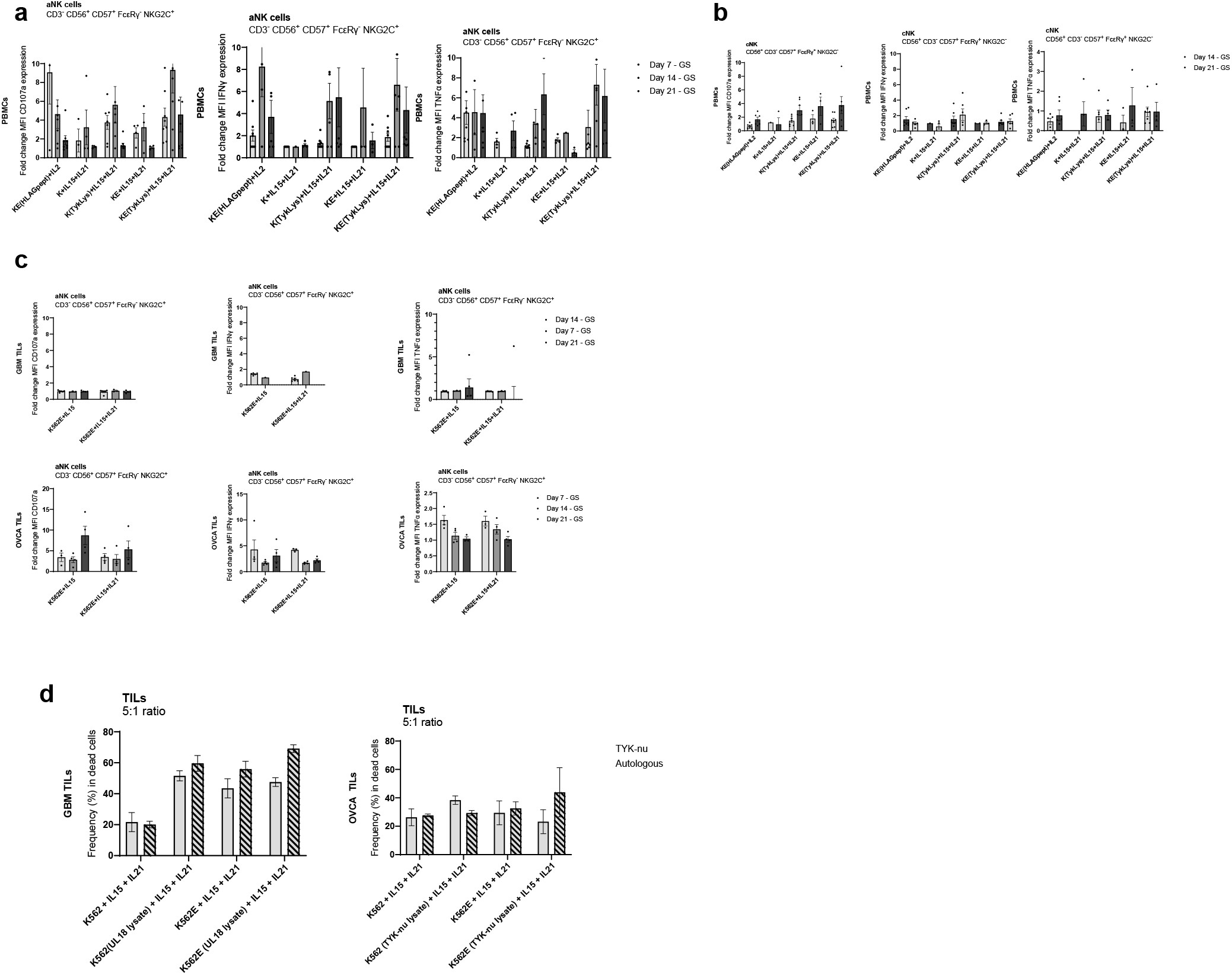
Functionality of aNK cells furing long-term expansion upon tumor antigen stimulation. **a**. Functional assay by flow cytometry performed on aNK cells on degranulation and cytokine production are studied with the MFI fold change, meaning MFI (stimulated sample) / MFI (unstimulated sample). PBMCs (*n=9*), GBM TILs (*n=6*) and IPLA-OVCA TILs (*n=4*) were generally stimulated with PMA/Ionomycin and specifically stimulated with viable UL18 for GBM TILs or TYK-nu for IPLA-OVCA TILs at 5:1 (E:T) ratio, and in all cases, cells were treated with anti-human CD107a antibody (PerCP/Cy5.5) and protein transport inhibitors Golgiplug and Golgistop. **b**. Functional assay by flow cytometry performed on cNK cells MFI fold change. **c**. Functional assay on TILs derived from GBM or OVCA patients. **d**. Killing assay of TILs derived from GBM or OVCA cells against general cell lines (UL18 for GBM and TYK-nu for OVCA) or autologous against the primary tumor cells. The positivity of the functionality was detemined with Fluorescence Minus One (FMO) tests. Data is presented as mean ± standard error of the mean (SEM), and statistical differences were tested using two-way ANOVA; *p<0,05, **p<0,01, ***p<0,001; ****p<0,0001.

### IL-21 and K562E TILs cultured with tumor lysate decreased tumor size in zebrafish model

After long-term expansion GBM TILs with IL-15 + IL-21 and feeder cells loaded with lysate, these cultures were transplanted into a zebrafish that was developing glioblastoma in the brain (**Fig. 6a**). We observed by confocal microscopy that tumor cells (in red) and immune cells (in green) translocated at the same places in the zebrafish embryos (**Fig. 6b**). When analyzing the tumor size, we observed that the samples that contained IL15 + IL-21 with K562E feeder cells that were loaded previously with the lysate were the ones that had the lowest tumor size after 3 days of transplantation, compared to the other controls (**Fig. 6c**).

## Discussion

Previous research has shown the effectiveness of expanding NK cells for therapeutic use with IL-15, particularly through genetic modifications and combinations with other cytokines like IL-21 [44]. This study introduces a novel protocol for expanding aNK cells by using a combination of IL-15 and IL-21, paired with K562E feeder cells. The approach offers a promising alternative to traditional methods by significantly enhancing NK cell expansion and functionality. Other studies have already proven the effectiveness of using IL-15 and IL-21 to expand NK cells [45]. Here, we show that the use of IL-21, in particular, boosts aNK cell proliferation and tumor cytotoxicity, which is critical for targeting solid tumors like glioblastoma and ovarian cancer. The importance of IL-15 and IL-21 combination has already been proven to expand NK cells and maintain high purity and viability [46]. This was proven successful in cancers like rhabdomyosarcoma. Here, we reinforce these findings, and we show that aNK cells are the predominant population increased when NK cells expand, proving their potential to target and kill tumors such as GBM and ovarian cancer.

Key results showed that the IL-15+IL-21 combination substantially increased NK cell frequency, achieving up to 80% NK cells by the end of the expansion without prior bead isolation. The protocol led to 14,000-fold NK cell expansion, with a selective 450-fold increase in aNK cells, demonstrating its robustness and efficiency. This selective expansion occurred without compromising the purity of the NK cells, as there was minimal T-cell contamination.

Functionally, while IL-21 contributed to the expansion of aNK cells, it did not significantly enhance their cytotoxicity or cytokine production beyond what was observed with IL-15 alone. Previous labs have already shown that IL-21 drives the functional maturation and expansion of NK cells, but it does not necessarily enhance their cytotoxic capabilities when used in combination with IL-15 [47]. This suggests that IL-21 mainly supports the proliferation of aNK cells rather than boosting their direct tumor-killing capabilities. Despite this, aNK cells exhibited sustained degranulation and cytokine production, albeit with a reduction over time, indicating the potential need for antigen-specific stimulation to maintain their long-term functionality.

Previous studies on NK cells derived from PBMCs have demonstrated higher cytotoxicity and persistence when expanded using feeder cells-free protocols, shown in ovarian cancer models. These expanded NK cells were more effective in killing tumor cells, particularly in solid tumors and metastatic tumors [48]. Similarly, in GBM, NK cells, although they infiltrate the TME, they are immunosuppressed. *Ex vivo* activation with cytokines (IL-15) and tumor lysates restore their cytotoxicity against GBM cells [49, 50]. In this paper, experiments using tumor lysates from glioblastoma and ovarian cancer revealed that aNK cells expanded under these conditions and displayed enhanced cytotoxicity against autologous tumor cells. This effect was more pronounced in aNK derived from PBMCs compared to aNK cells derived from TILs, suggesting that circulating NK cells may respond differently to the TME. NK cells derived from PBMCs maintain their natural receptor profiles (NKG2D), which is crucial for recognizing and eliminating tumor cells [50]. The study also showed that aNK cells maintained their natural receptor expression profiles, including the upregulation of activating receptors like NKG2D, enhancing their anti-tumor function. These findings suggest that circulating NK cells, especially when expanded and activated outside the immunosuppressive TME, display more robust anti-tumor activity. This highlights the importance of NK cell expansion and receptor profile maintenance for effective cancer immunotherapy.

*In vivo* studies using a zebrafish model further supported the therapeutic potential of this protocol. Previous studies showed the expansion of NK cells using K562 feeder cells expressing OX40 ligand, which resulted in enhanced NK cell proliferation and sustained functionality [34, 35]. These papers reinforce our study, in which TILs expanded with IL-15+IL-21 and K562E feeder cells, pre-loaded with tumor lysates, demonstrated significant anti-tumor activity. These cells successfully migrated to tumor sites, resulting in reduced tumor size in the zebrafish embryos, highlighting the potential of this method for improving TIL-based therapies.

In conclusion, this study presents an effective protocol for expanding aNK cells using IL-15 and IL-21, which enhances both their proliferation and their ability to target tumors. The approach holds promise for advancing aNK cell-based therapies, particularly in cold tumors such as glioblastoma and ovarian cancer. Future research should focus on optimizing antigen stimulation to sustain long-term proliferation and fully realize the therapeutic potential of aNK cells.

## REFERENCES

1. Park, J.H., et al., Long-Term Follow-up of CD19 CAR Therapy in Acute Lymphoblastic Leukemia. N Engl J Med, 2018. 378(5): p. 449–459.

2. Gill, S., M.V. Maus, and D.L. Porter, Chimeric antigen receptor T cell therapy: 25years in the making. Blood Rev, 2016. 30(3): p. 157–67.

3. Fry, T.J., et al., CD22-targeted CAR T cells induce remission in B-ALL that is naive or resistant to CD19-targeted CAR immunotherapy. Nat Med, 2018. 24(1): p. 20–28.

4. Melenhorst, J.J., et al., Decade-long leukaemia remissions with persistence of CD4(+) CAR T cells. Nature, 2022. 602(7897): p. 503–509.

5. Safarzadeh Kozani, P., et al., Recent Advances in Solid Tumor CAR-T Cell Therapy: Driving Tumor Cells From Hero to Zero? Front Immunol, 2022. 13: p. 795164.

6. Newick, K., et al., CAR T Cell Therapy for Solid Tumors. Annu Rev Med, 2017. 68: p. 139–152.

7. Laskowski, T.J., A. Biederstädt, and K. Rezvani, Natural killer cells in antitumour adoptive cell immunotherapy. Nat Rev Cancer, 2022. 22(10): p. 557–575.

8. Mujal, A.M., R.B. Delconte, and J.C. Sun, Natural Killer Cells: From Innate to Adaptive Features. Annu Rev Immunol, 2021. 39: p. 417–447.

9. Shimasaki, N., A. Jain, and D. Campana, NK cells for cancer immunotherapy. Nat Rev Drug Discov, 2020. 19(3): p. 200–218.

10. Raskov, H., et al., Natural Killer Cells in Cancer and Cancer Immunotherapy. Cancer Lett, 2021. 520: p. 233–242.

11. Lanier, L.L., NK cell recognition. Annu Rev Immunol, 2005. 23: p. 225–74.

12. Orr, M.T. and L.L. Lanier, Natural killer cell education and tolerance. Cell, 2010. 142(6): p. 847–56.

13. Ljunggren, H.G. and K. Kärre, In search of the ‘missing self’: MHC molecules and NK cell recognition. Immunol Today, 1990. 11(7): p. 237–44.

14. Shenk, T. and J.C. Alwine, Human Cytomegalovirus: Coordinating Cellular Stress, Signaling, and Metabolic Pathways. Annu Rev Virol, 2014. 1(1): p. 355–74.

15. Hammer, Q., et al., Peptide-specific recognition of human cytomegalovirus strains controls adaptive natural killer cells. Nat Immunol, 2018. 19(5): p. 453–463.

16. López-Botet, M., et al., Adaptive NK cell response to human cytomegalovirus: Facts and open issues. Semin Immunol, 2023. 65: p. 101706.

17. Cooper, M.A., et al., Cytokine-induced memory-like natural killer cells. Proc Natl Acad Sci U S A, 2009. 106(6): p. 1915–9.

18. Romee, R., et al., Cytokine activation induces human memory-like NK cells. Blood, 2012. 120(24): p. 4751–60.

19. Terrén, I., et al., Cytokine-Induced Memory-Like NK Cells: From the Basics to Clinical Applications. Front Immunol, 2022. 13: p. 884648.

20. Sarhan, D., et al., Adaptive NK Cells with Low TIGIT Expression Are Inherently Resistant to Myeloid-Derived Suppressor Cells. Cancer Res, 2016. 76(19): p. 5696–5706.

21. Sarhan, D., et al., Adaptive NK Cells Resist Regulatory T-cell Suppression Driven by IL37. Cancer Immunol Res, 2018. 6(7): p. 766–775.

22. Kamiya, T., et al., Blocking expression of inhibitory receptor NKG2A overcomes tumor resistance to NK cells. J Clin Invest, 2019. 129(5): p. 2094–2106.

23. Nikzad, R., et al., Human natural killer cells mediate adaptive immunity to viral antigens. Sci Immunol, 2019. 4(35).

24. Stary, V., et al., A discrete subset of epigenetically primed human NK cells mediates antigen-specific immune responses. Sci Immunol, 2020. 5(52).

25. Siemaszko, J., A. Marzec-Przyszlak, and K. Bogunia-Kubik, Activating NKG2C Receptor: Functional Characteristics and Current Strategies in Clinical Applications. Arch Immunol Ther Exp (Warsz), 2023. 71(1): p. 9.

26. Bryceson, Y.T., et al., Synergy among receptors on resting NK cells for the activation of natural cytotoxicity and cytokine secretion. Blood, 2006. 107(1): p. 159–66.

27. Fujisaki, H., et al., Expansion of highly cytotoxic human natural killer cells for cancer cell therapy. Cancer Res, 2009. 69(9): p. 4010–7.

28. Miller, J.S., et al., A First-in-Human Phase I Study of Subcutaneous Outpatient Recombinant Human IL15 (rhIL15) in Adults with Advanced Solid Tumors. Clin Cancer Res, 2018. 24(7): p. 1525–1535.

29. Skrombolas, D. and J.G. Frelinger, Challenges and developing solutions for increasing the benefits of IL-2 treatment in tumor therapy. Expert Rev Clin Immunol, 2014. 10(2): p. 207–17.

30. Knorr, D.A., et al., Clinical utility of natural killer cells in cancer therapy and transplantation. Semin Immunol, 2014. 26(2): p. 161–72.

31. Cho, D. and D. Campana, Expansion and activation of natural killer cells for cancer immunotherapy. Korean J Lab Med, 2009. 29(2): p. 89–96.

32. Li, Q., et al., Multiple effects of IL-21 on human NK cells in ex vivo expansion. Immunobiology, 2015. 220(7): p. 876–88.

33. Granzin, M., et al., Highly efficient IL-21 and feeder cell-driven ex vivo expansion of human NK cells with therapeutic activity in a xenograft mouse model of melanoma. Oncoimmunology, 2016. 5(9): p. e1219007.

34. Kweon, S., et al., Expansion of Human NK Cells Using K562 Cells Expressing OX40 Ligand and Short Exposure to IL-21. Front Immunol, 2019. 10: p. 879.

35. Yano, M., et al., Evaluation of allogeneic and autologous membrane-bound IL-21-expanded NK cells for chronic lymphocytic leukemia therapy. Blood Adv, 2022. 6(20): p. 5641–5654.

36. Leong, J.W., et al., Preactivation with IL-12, IL-15, and IL-18 induces CD25 and a functional high-affinity IL-2 receptor on human cytokine-induced memory-like natural killer cells. Biol Blood Marrow Transplant, 2014. 20(4): p. 463–73.

37. Liu, L.L., et al., Ex Vivo Expanded Adaptive NK Cells Effectively Kill Primary Acute Lymphoblastic Leukemia Cells. Cancer Immunol Res, 2017. 5(8): p. 654–665.

38. Phan, M.T., et al., Selective Expansion of NKG2C+ Adaptive NK Cells Using K562 Cells Expressing HLA-E. Int J Mol Sci, 2022. 23(16).

39. Haroun-Izquierdo, A., et al., Adaptive single-KIR(+)NKG2C(+) NK cells expanded from select superdonors show potent missing-self reactivity and efficiently control HLA-mismatched acute myeloid leukemia. J Immunother Cancer, 2022. 10(11).

40. Yang, H., et al., Visualization of natural killer cell-mediated killing of cancer cells at single-cell resolution in live zebrafish. Biosens Bioelectron, 2022. 216: p. 114616.

41. Murali Shankar, N., et al., Preclinical assessment of CAR-NK cell-mediated killing efficacy and pharmacokinetics in a rapid zebrafish xenograft model of metastatic breast cancer. Front Immunol, 2023. 14: p. 1254821.

42. Pascoal, S., et al., A Preclinical Embryonic Zebrafish Xenograft Model to Investigate CAR T Cells In Vivo. Cancers (Basel), 2020. 12(3).

43. Zhu, Z., et al., A novel cuproptosis-related molecular pattern and its tumor microenvironment characterization in colorectal cancer. Front Immunol, 2022. 13: p. 940774.

44. Islam, R., et al., Enhancing a Natural Killer: Modification of NK Cells for Cancer Immunotherapy. Cells, 2021. 10(5).

45. Zhang, C., et al., Sequential Exposure to IL21 and IL15 During Human Natural Killer Cell Expansion Optimizes Yield and Function. Cancer Immunol Res, 2023. 11(11): p. 1524–1537.

46. Wagner, J., et al., A Two-Phase Expansion Protocol Combining Interleukin (IL)-15 and IL-21 Improves Natural Killer Cell Proliferation and Cytotoxicity against Rhabdomyosarcoma. Front Immunol, 2017. 8: p. 676.

47. Brady, J., et al., IL-21 induces the functional maturation of murine NK cells. J Immunol, 2004. 172(4): p. 2048–58.

48. Chen, M., et al., Anti-Tumor Activity of Expanded PBMC-Derived NK Cells by Feeder-Free Protocol in Ovarian Cancer. Cancers (Basel), 2021. 13(22).

49. Pan, C., et al., NK Cell-Based Immunotherapy and Therapeutic Perspective in Gliomas. Front Oncol, 2021. 11: p. 751183.

50. Burger, M.C., et al., CAR-Engineered NK Cells for the Treatment of Glioblastoma: Turning Innate Effectors Into Precision Tools for Cancer Immunotherapy. Front Immunol, 2019. 10: p. 2683.

